# Opening the Black Box of Neural Computation from Neural Recordings with Gain-Modulated Linear Dynamical System

**DOI:** 10.64898/2026.03.12.711252

**Authors:** Yiteng Zhang, Zixiong Wang, Xingyu Li, Bin Min

**Affiliations:** Lingang Laboratory, Shanghai, China; Institute of Neuroscience, Key Laboratory of Brain Cognition and Brain-Inspired Intelligence Technology, CAS Center for Excellence in Brain Science and Intelligence Technology, Chinese Academy of Sciences, Shanghai 200031, China; University of Chinese Academy of Sciences, Beijing 100049, China

## Abstract

Inferring computational mechanisms from neural recordings is a central goal in systems neuroscience. Recent developments have identified low-rank recurrent neural networks (RNNs) as an effective tool for fitting observed neural activity and extracting neural dynamics. However, we show that accurate activity fitting alone does not guarantee mechanistic validity: even well-fitted low-rank RNNs can yield misleading circuit interpretations in the absence of ground-truth model settings on synthetic datasets. To address this limitation, we introduce a gain-modulated linear dynamical systems (gmLDS) method, which decomposes latent-variable interactions into a time-varying gain and static low-rank connectivity components. This decomposition captures linearized dynamics near neural trajectories and enables flexible adaptation to each neuron’s nonlinear responses without requiring a predefined activation function. Extensive validation across multiple synthetic datasets shows that gmLDS accurately reproduces neural activity, gain and connectivity, thereby capturing the fine structure of linearized dynamics and the underlying circuit mechanisms. When applying to neural recordings from a context-dependent decision-making task, gmLDS uncovers evidence for the coexistence of two prevalent selection mechanisms, offering new insights into a long-standing unresolved issue in the field. Together, our results establish gmLDS as a principled approach for opening the black box of neural computation from neural recordings.

## Introduction

Inferring the computational mechanisms underlying neural population activity is a fundamental challenge in systems neuroscience (Romo et al. 1999; Mante et al. 2013; Cunningham et al. 2014; Semedo et al. 2019; Aoi et al. 2020). While recent advances in machine learning have provided powerful tools for fitting large-scale neural recordings (Linderman et al. 2017; Schneider et al. 2023; Abbaspourazad et al. 2023; Nair et al. 2023; Langdon et al. 2025), a significant gap remains: high predictive accuracy does not necessarily translate to mechanistic insight. Most current latent-variable models function as “black boxes”, offering little transparency into how connectivity and dynamics conspire to perform specific computations.

A promising framework for bridging this gap is the low-rank Recurrent Neural Network (RNN) (Mastrogiuseppe et al. 2018; Schuessler, Dubreuil, et al. 2020; Schuessler, Mastrogiuseppe, et al. 2020; Beiran et al. 2021; Valente et al. 2022; Valente et al. 2022; Dubreuil et al. 2022; Galgali et al. 2023; Beiran et al. 2023; Vinograd et al. 2024; Ostojic et al. 2024; Zhang et al. 2025). Theoretically, low-rank RNNs provide a transparent link between a network’s connectivity, its resulting state-space dynamics, and its underlying computation. This theoretical clarity has inspired new inference methods, such as Low-rank Inference from Neural Trajectories (LINT) (Valente et al. 2022), which attempt to extract circuit mechanisms directly from data. However, these methods are often “brittle”; they typically require the researcher to pre-define the neuron’s activation function (e.g., Tanh or ReLU). We show that when these pre-defined assumptions do not perfectly match the ground truth, inferred connectivity becomes biased, leading to incorrect conclusions about the underlying circuit mechanisms (see 4.1 for details).

To overcome these limitations, we introduce the gain-modulated Linear Dynamical Systems (gmLDS). Our approach reformulates the low-rank RNN framework into a flexible dynamical system by decomposing latent-variable interactions into two distinct components: a time-varying, state-dependent neural gain and a static, low-rank connectivity matrix. In fact, the nonlinear dynamics of a typical low-rank RNN can be described by the collection of linearized local dynamics at each time point along the network trajectory (see Fig. 1), where the local linearized dynamics are determined not only by the static connectivity but also the excitation level of neurons at corresponding time (Sussillo et al. 2013; Vyas et al. 2020; Pagan et al. 2024; Zhang et al. 2025). The neural gain term in our model captures this time-dependent excitation state with a Multi-Layer Perceptron (MLP). In this way, gmLDS flexibly adapts to a wide range of nonlinear responses without requiring a predefined activation function. Consequently, gmLDS yields a complete mechanistic description of neural computation by simultaneously inferring latent dynamics, time-varying gains, and the underlying connectivity.

**Figure 1:**
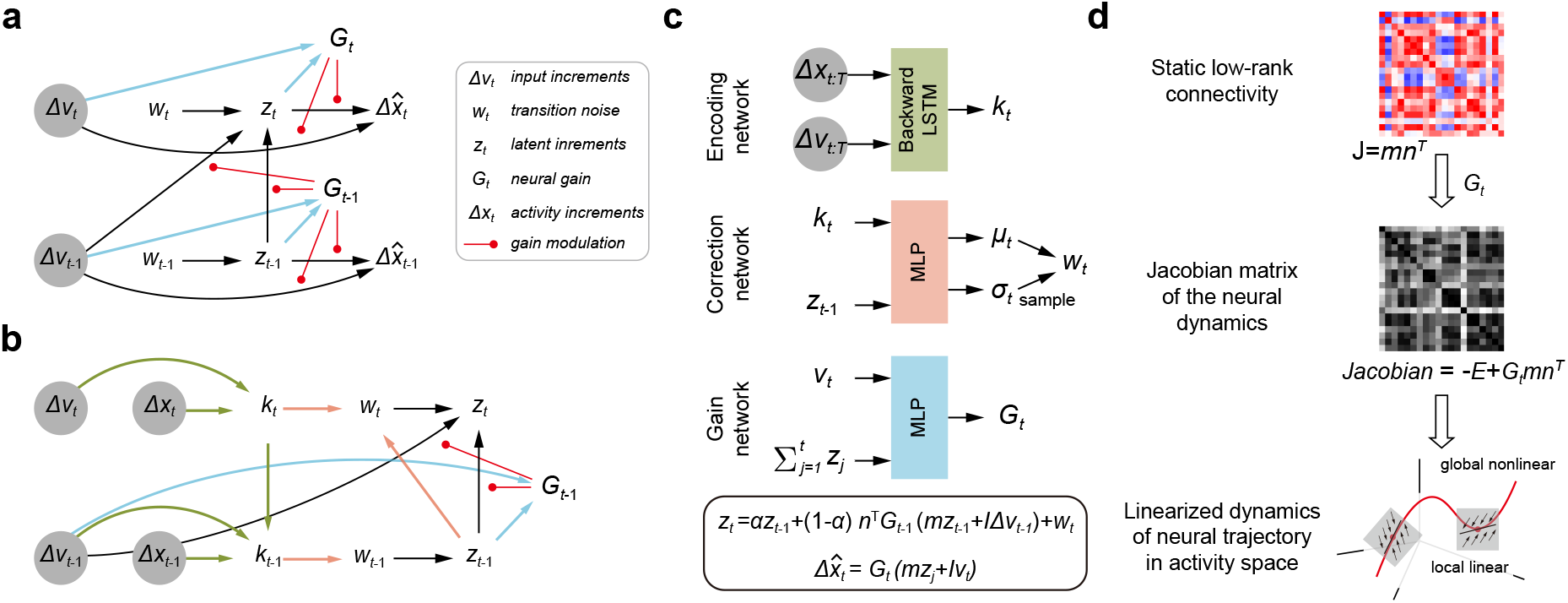
Graphical description of the gain-modulated latent dynamical system (gmLDS). **a**, The generative model of gmLDS. **b**, The posterior inference model of gmLDS. **c**, Detailed computational modules corresponding to panels a and b. A backward LSTM embeds future information to *k*_*t*_; a correction module infers the posterior correction term *w*_*t*_; and a gain module computes the time-varying neural gain *G*_*t*_. Δ*x*_*t*:*T*_ and Δ*ν*_*t*:*T*_ denote the sequence {Δ*x*_*t*_, …, Δ*x*_*T*_} and {Δ*ν*_*t*_, …, Δ*ν*_*T*_}, respectively. **d**, gmLDS infers a static low-rank connectivity and a time-varying gain, providing an estimate of the linearized dynamics along neural trajectories and enabling analysis of fixed points and line attractors.

The contributions of this work are three-fold:

1. Methodological Innovation: Grounded in low-rank RNN theory, we develop a data-driven approach that recovers connectivity and computation without making restrictive assumptions about the underlying neural activation functions.
2. Robust Validation: Through extensive testing on diverse synthetic datasets, we demonstrate that gmLDS outperforms existing methods in accurately recovering the “ground truth” connectivity and linearized dynamics.
3. Biological Discovery: Applying gmLDS to neural recordings from a context-dependent decision-making (CDM) task, we find evidence for the coexistence of two prevalent selection mechanisms (Pagan et al. 2024). This result offers a potential reconciliation for a long-standing debate in the field (Mante et al. 2013; Valente et al. 2022; Soldado-Magraner et al. 2024; Zhang et al. 2025) regarding how the brain filters task-irrelevant information.

Together, our results establish gmLDS as a principled framework for opening the “black box” of neural computation from neural recordings. By decoupling gain from connectivity, we provide a robust path toward identifying the circuit-level implementations of cognitive functions.

## Methods

The gain-modulated latent dynamical system (gmLDS) is implemented in a manner similar to the variational RNN (Chung et al. 2015), including a **generative model** that specifies the temporal dynamics of the system and a **posterior inference model** that enables parameter updating in the generative model through variational inference. We dive into implementation details in the following subsections.

### Generative model

We consider trials of length *T* + 1 with neural population activity 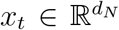 and stimulus input 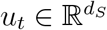. Neural activity increments are defined by

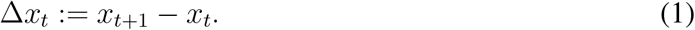

The stimulus input *u* was low-pass filtered with time constant *τ*,

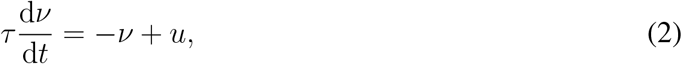

and define the corresponding input increments as Δ*ν*_*t*_ := *ν*_*t*+1_ − *ν*_*t*_. Under this formulation, the generative process in the differential domain is defined as:

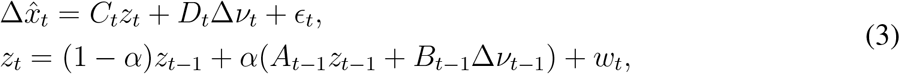

where 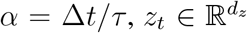 denotes increments in latent space, *ϵ*_*t*_ ∼ 𝒩 (0, *Q*) and *w*_*t*_ ∼ 𝒩 (0, *R*) denote observation noise and transition noise sampled from prior distribution, respectively.

The time-varying linear operators *A*_*t*_, *B*_*t*_, *C*_*t*_, *D*_*t*_ are defined through a shared gain-modulated factorization:

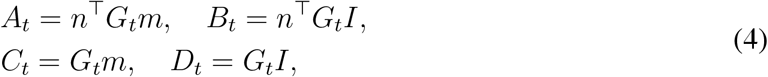

where *G*_*t*_ = GainNetwor 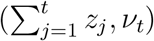 (blue arrow in Fig. 1a, b) is a diagonal gain matrix computed from the accumulated latent state and relative integrated stimulus input via a multilayer perceptron. Matrices *m* and *n* encode static low-rank recurrent connectivity, while *I* represents static input connectivity.

Together, these equations define a gain-modulated latent dynamical system in the differential domain. Given stimulus input, the model specifies how local latent increments generate neural activity increments through state-dependent linear operators, and how latent increments evolve over time. The observed neural trajectory is obtained by cumulatively summing the generated activity increments.

### Posterior inference model

The goal of posterior inference in gmLDS is to estimate the smoothed posterior distribution over latent increments, *p*(*z*_*t*_|Δ*x*_1:*T*_, Δ*ν*_1:*T*_), given neural activity increments and stimulus input increments over an entire trial. Direct inference of this posterior is analytically intractable, because both the latent transition and observation models depend nonlinearly on the latent state. This nonlinear dependence breaks the linear–Gaussian structure required for closed-form Bayesian inference.

To make inference tractable, we exploit the Markov structure of the latent dynamics and rewrite the posterior as

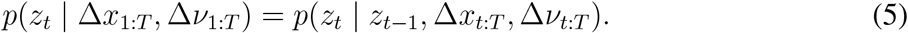

We introduce a context variable *k*_*t*_ to summarize information from future observations, allowing the posterior to be expressed compactly as *p*(*z*_*t*_ | *z*_*t*−1_, *k*_*t*_).

Rather than modeling this posterior directly over *z*_*t*_, we place stochasticity in the transition noise driving the latent dynamics. Specifically, the latent update is written as

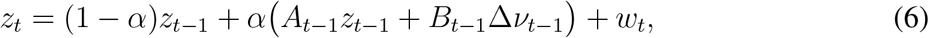

where uncertainty in the posterior is captured by the transition noise *w*_*t*_. In contrast to the generative model in Eq. (3), the transition noise is here sampled from a learned posterior distribution defined by the correction model (orange arrow in Fig. 1b),

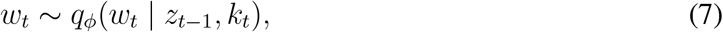

where *q*_*ϕ*_ is a multivariate Gaussian whose mean and covariance are parameterized by a MLP. By placing stochasticity in the transition noise rather than in the latent state itself, the latent dynamics evolve deterministically once the noise is sampled.

To construct the context variables {*k*_*t*_}, we encode neural activity increments and stimulus input increments from time step *t* to *T* using a backward LSTM (green arrow in Fig. 1b) (Hochreiter et al. 1997),

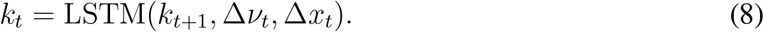

### Learning and optimization

Model parameters are learned by maximizing a variational evidence lower bound (ELBO) on the marginal log-likelihood of neural activity. Let *θ* denote the parameters of the generative model and *ϕ* denote those of the correction model. Given the observed sequences of neural activity increments {Δ*x*_*t*_} and stimulus input increments {Δ*ν*_*t*_}, we optimize the following negative ELBO:

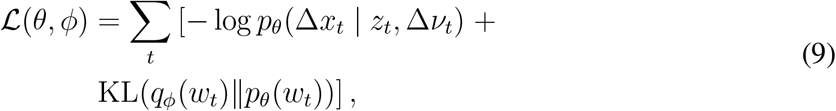

where KL denotes the KL-divergence between the posterior and the prior distribution.

The reconstruction term log *p*_*θ*_(Δ*x*_*t*_|*z*_*t*_, Δ*ν*_*t*_) encourages the generative model to accurately reproduce observed neural activity increments, while the KL divergence regularizes the inferred transition noise toward its prior distribution.

All model parameters—including the static connectivity matrices (*m, n, I*), the prior covariance *R* and *Q* in the generative model, the gain network, the posterior encoding and correction network —are optimized jointly via gradient descend. To enable optimization through the stochastic latent transitions, we employ the reparameterization trick (Kingma et al. 2022) to compute gradients through the sampled transition noise.

### Relationship to the low-rank RNN

A central motivation for gmLDS is to provide direct, data-driven evaluation of the local linearized dynamics that have long been used to interpret computation in RNNs (Sussillo et al. 2013). Classical linearization-based analyses require full access to the trained model, including both the recurrent connectivity and the neuronal activation functions. In biological neural circuits, neither quantity is directly observable, making it difficult to recover local linearized dynamics solely from neural recordings. Here, we show that gmLDS enables decent fitting to the local linearized dynamics directly from the observed neural activity without prior knowledge of connectivity and activation functions.

Consider a low-rank RNN of rank-*r*:

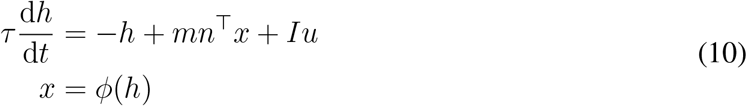

where *τ* is time constant, 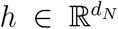 represents the neural activation, 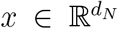 denotes neural activity, *ϕ*(·) is the nonlinear activation function, and the matrices *m*, 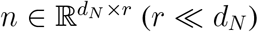 define the low-rank recurrent structure.

In such networks, the neural activation is confined in the space spanned by *m* and *I*, and can be writen as a linear combination of *m* and *I* as *h* = *mκ* + *Iν*. The latent dynamics follows:

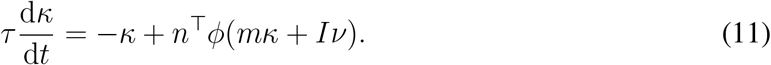

After discretizations of the above equation using Euler’s method with time step Δ*t*, and linearizing *ϕ*(·) around *h*_*t*_, the dynamics of the latent increments Δ*κ*_*t*_ = *κ*_*t*+1_ − *κ*_*t*_ reads

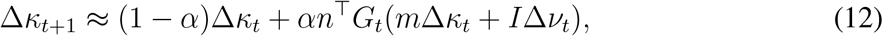

where *α* = Δ*t/τ* and *G*_*t*_ = diag(*ϕ*^′^(*h*_*t*_)) being the gain matrix. Furthermore, the differential neural activity follows the linearized readout:

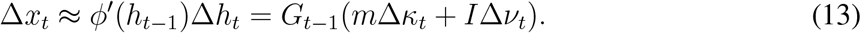

revealing that the proposed system effectively acts as a locally linear surrogate for low-rank RNNs in the differential domain.

More importantly, the local linearization of a low-rank RNN around a reference activity state *x*_0_ = *ϕ*(*h*_0_) takes the form:

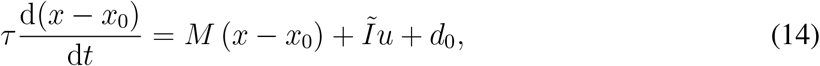

where *Ĩ* = *G*(*h*_0_)*I* denotes input representation direction, *d*_0_ denotes the instantaneous velocity evaluated at *x*_0_, and *M* denotes the Jacobian matrix of the dynamical system, given by

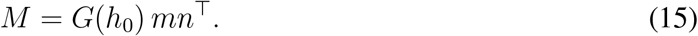

Here, *G*(*h*_0_) = diag(*ϕ*^′^(*h*_0_)) is the neuronal gain matrix evaluated at the linearization point. This implies that once gmLDS has inferred the time-varying gain *G*_*t*_ together with the static connectivity factors *m* and *n*, it directly provides an estimate of the transient matrix governing the local linearized dynamics along the neural trajectory.

## Experimental Results

### gmLDS recovers ground-truth gain and connectivity across diverse cognitive tasks

To evaluate gmLDS under controlled conditions spanning diverse cognitive tasks, we applied the model to synthetic neural population activity generated from RNNs. These ground-truth RNNs were trained to perform a range of classical cognitive tasks, including Perceptual Decision Making (PDM), Contextual Decision Making (CDM), Parametric Working Memory (PWM), and Delayed Match to Sample (DMS). This setup provides a principled testbed in which both neural activity and the underlying computational mechanisms are known, enabling quantitative assessment of a model’s ability to recover latent dynamics, gain modulation, and connectivity structure.

Our results demonstrate that all evaluated models are capable of accurately reconstructing neural population activity (Fig. 2a top, 2b; Supp. Table. 1, *R*^2^ of activity fitting). However, the LINT baseline failed to correctly identify the neural gain—defined as the derivative of the activation function—due to the imposed nonlinearity mismatch (Fig. 2a bottom, 2c; Supp. Table. 1, *R*^2^ of gain inferring). In contrast, gmLDS successfully inferred the ground-truth neural gain profiles across all tasks and RNN configurations. This underscores the effectiveness of gmLDS in capturing the linearized dynamics of the system.

**Figure 2:**
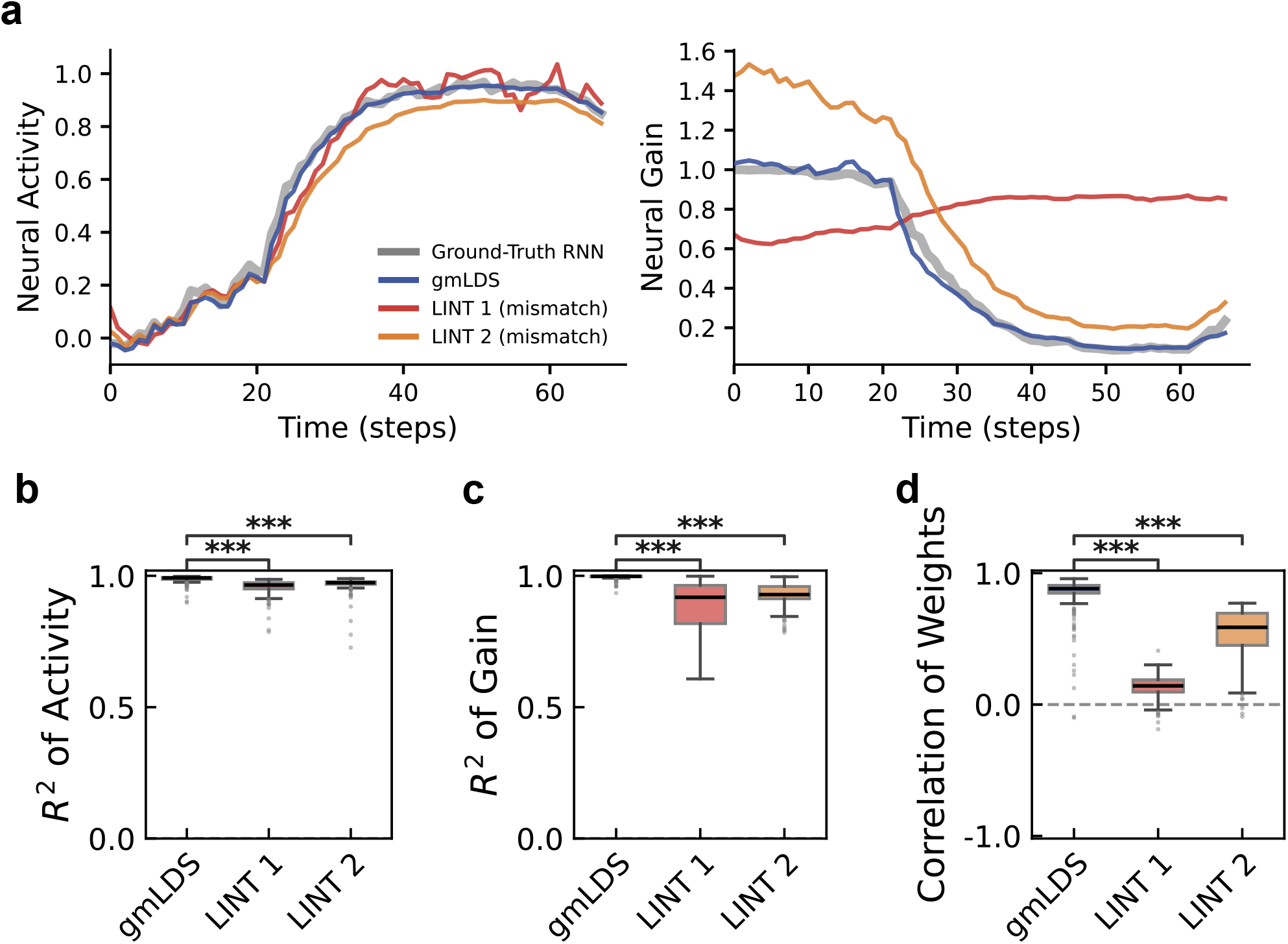
gmLDS accurately recovers gain and connectivity in CDM task. **a**, Example trajectories of neural activity (left) and neural gain (right) for a single trial. **b**, Reconstruction accuracy of neural population activity (*R*^2^). **c**, Inference accuracy of state-dependent neural gain (*R*^2^). **d**, Correlation between inferred and ground-truth recurrent connectivity weights. Boxplots show median, quartiles, and whiskers (1.5 × IQR). Asterisks denote statistical significance (*** *p <* 0.001, Wilcoxon signed-rank test). LINT 1, LINT 2, and the ground-truth RNN use three different activation functions.

Furthermore, we examined the performance of gmLDS in connectivity inference. By computing the correlation between the ground-truth and inferred recurrent synaptic inputs for individual neurons, we verified that the connectivity matrices recovered by gmLDS successfully match the ground-truth values (Fig. 2d; Supp. Table. 1, Correlation of connectivity). By integrating the inferred static connectivity structure with the state-dependent neural gains during task performance, gmLDS provides a robust framework for deciphering the neural substrates of computation.

### gmLDS can accurately infer computational mechanism in synthetic neural data

A central question is whether local linearized dynamics and the computational mechanisms they imply can be reliably inferred from neural activity alone. To address this, we first apply gmLDS to the CDM task, a canonical paradigm in which the computational role of linearized dynamics has been extensively characterized (Mante et al. 2013; Pagan et al. 2024; Zhang et al. 2025). In the CDM task, subjects must selectively accumulate the task-relevant sensory evidence while suppressing irrelevant information based on context. Previous modeling studies have shown that RNNs trained to perform this task exhibit dynamics that are well approximated by a linear attractor near the choice axis in each context (Mante et al. 2013). In the absence of external input, the dynamics can be described by a linear system, 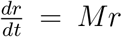, where the Jacobian matrix *M* has a single eigenvalue close to 0 and all remaining eigenvalues having negative real parts. The left eigenvector associated with the zero eigenvalue defines a direction in neural activity space along which activity is preserved over time, while activity along all other directions will decay to zero. As a consequence, only the component of a input representation (***Ĩ***) that projects onto this direction can be integrated to influence the decision variable. This vector therefore determines which aspects of the input are selected for accumulation and is referred to as the **selection vector *s*** (Mante et al. 2013).

Given the concept of selection vector, two distinct selection mechanism is introduced (Pagan et al. 2024) (Fig. 3b). Because the contribution of a sensory input to evidence accumulation is determined by its projection ***Ĩ***·***s***, differences between relevant and irrelevant contexts can therefore result either from changes in the input representation or from changes in the selection vector itself. These two contributions can be formally decomposed as :

**Figure 3:**
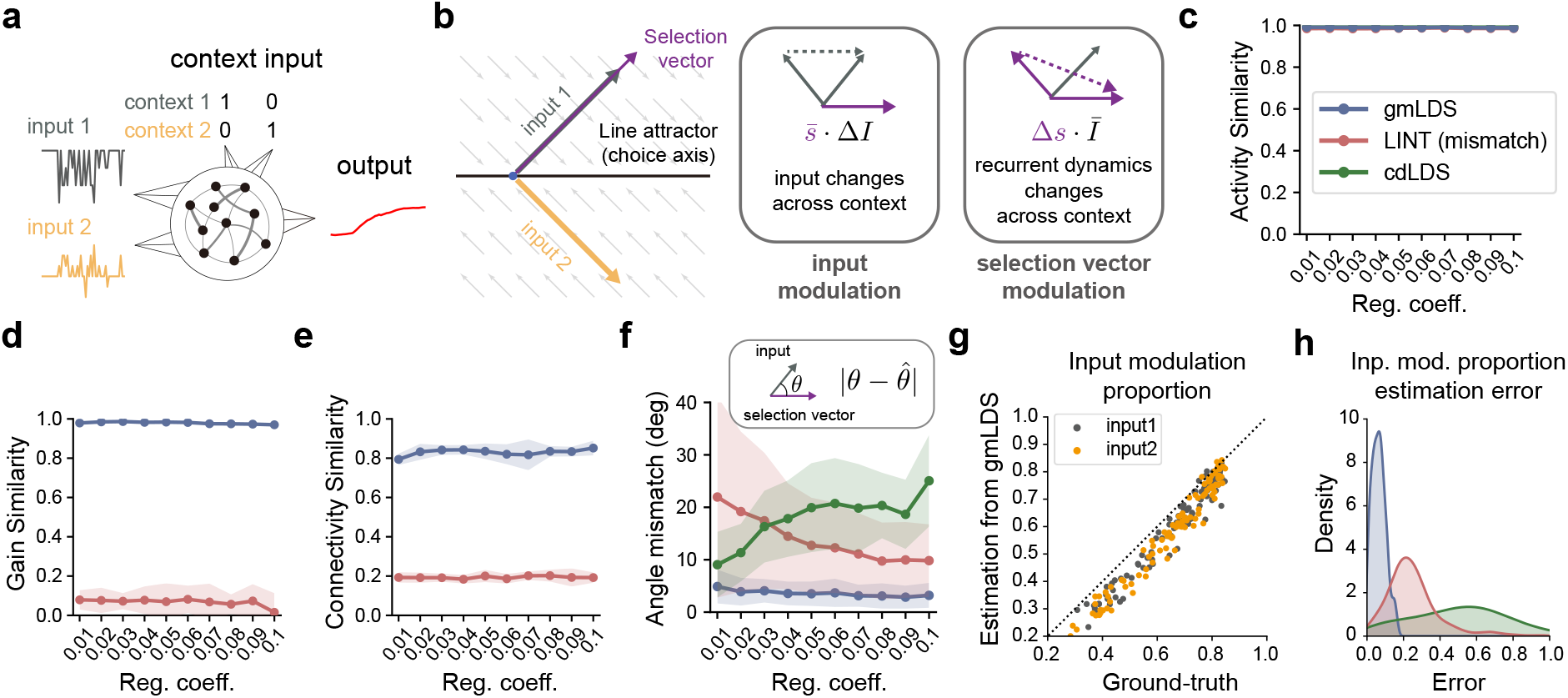
gmLDS recovers the geometry and selection mechanism of context-dependent computation in synthetic data. **a**, Experimental setting of vanilla RNNs trained to perform the context-dependent decision-making (CDM) task. **b**, Two candidate mechanisms for CDM: input modulation and selection vector modulation. **c**, Similarity between inferred and ground-truth neural activity across methods. Reg. coeff., regulization coefficient. cdLDS stands for context-dependent LDS (Soldado-Magraner et al. 2024). **d**, Similarity between inferred and ground-truth neuronal gain. **e**, Similarity between inferred and ground-truth low-rank connectivity. **f**, Error between inferred and ground-truth angles. **g**, Estimated versus ground-truth input modulation strength. **h**, Distribution of absolute estimation errors across methods. Inp. mod. denotes input modulation.

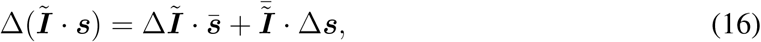

where Δ denotes the difference between contexts and the overbar denotes the context average. The first term corresponds to input modulation, whereas the second term corresponds to selection vector modulation. Despite the conceptual clarity of this decomposition, there is currently no direct way to estimate either term from neural recordings alone, motivating the application of gmLDS to the CDM task.

To evaluate gmLDS in a controlling setting, we generate synthetic data using vanilla RNNs trained on a CDM task (Fig. 3a, See methods for more details). To systematically manipulate the underlying selection mechanism, we varied the strength of regularization applied to the recurrent connectivity during training. Larger values of *λ* encourage lower-dimensional neural activities, biasing network toward input modulation, whereas weaker regularization permitted higher-dimensional activities that support selection vector modulation (Zhang et al. 2025). For each regularization strength, we trained 10 independently initialized RNNs for subsequent analysis.

We first asked whether gmLDS can accurately reconstruct neural activity itself. As shown in Fig. 3c, gmLDS achieves high activity-level similarity to the ground-truth RNN across a wide range of regularization strengths, as quantified by the average *R*^2^ score computed across neurons. Importantly, this level of activity reconstruction is comparable to that obtained by baseline methods, including LINT and a context-dependent LDS (cdLDS) approach that fits separate models for each context (Soldado-Magraner et al. 2024). Notably, LINT assumes a activation function that differs from that of the ground-truth RNN, yet all methods achieve similarly high activity-level reconstruction performace.

However, accurate activity reconstruction alone does not guarantee mechanistic fidelity. As shown in Fig. 2, models with similar activity-level fits can differ substantially in their inferred mechanisms. We therefore examined whether the inferred models recover the underlying linearized dynamics by comparing their estimates of neuronal gain and recurrent connectivity with the ground-truth values. As shown in Fig. 3d,e, gmLDS shows strong correspondence with the ground-truth gain (Fig. 3d) and more accurately recovers the structure of the recurrent connectivity than LINT (Fig. 3e). The cdLDS baseline does not explicitly parameterize gain or connectivity at the single-neuron level, and is therefore not included in these comparisons.

Beyond recovering parameters, we next assessed whether gmLDS preserves the geometry underlying selection. Specifically, we quantified the angle between the inferred selection vector and the sensory input direction and compared it with the ground-truth value. As shown in Fig. 3f, gmLDS yields substantially smaller angular errors, indicating more accurate recovery of the geometric relationships between ***Ĩ*** and ***s*** that determine which input is integrated by the line attractor. For the cdLDS model, the input directions and selection vectors are defined in the latent space and subsequently projected into the neural activity space via the observation matrix before computing the corresponding angles.

Having established that gmLDS accurately recovers the linearized dynamics of trained CDM RNNs, we finally asked whether it enables quantitative inference of the underlying selection mechanisms. Using the inferred selection vectors and input representations, we computed the contributions of input modulation and selection vector modulation according to Eq. (16). Fig. 3g shows that gmLDS-estimated input modulation closely matches the ground-truth value across networks, with points tightly aligned along the identity line. To further quantify estimation accuracy, we compared the absolute errors in inferred input modulation across methods. Consistent with this observation, the distribution of absolute estimation errors (Fig. 3h) shows that gmLDS yields significantly smaller errors than baseline approaches. Together, these results demonstrate that gmLDS can accurately infer underlying computational mechanisms from synthetic neural data, as illustrated by the CDM example.

### gmLDS unveils line-attractor dynamics and context-dependent modulation in primate PFC recordings

To test whether gmLDS can uncover circuit-level computational mechanisms from real neural recordings, we applied our method to *in vivo* electrophysiological data from macaque monkeys performing a CDM task (Mante et al. 2013) (Fig. 4a). In this task, animals viewed a random-dot stimulus conveying evidence along two sensory dimensions—motion direction and dot color—while a contextual cue instructed them to report either motion or color on each trial. Neural activity was recorded from 727 neurons in the prefrontal cortex.

**Figure 4:**
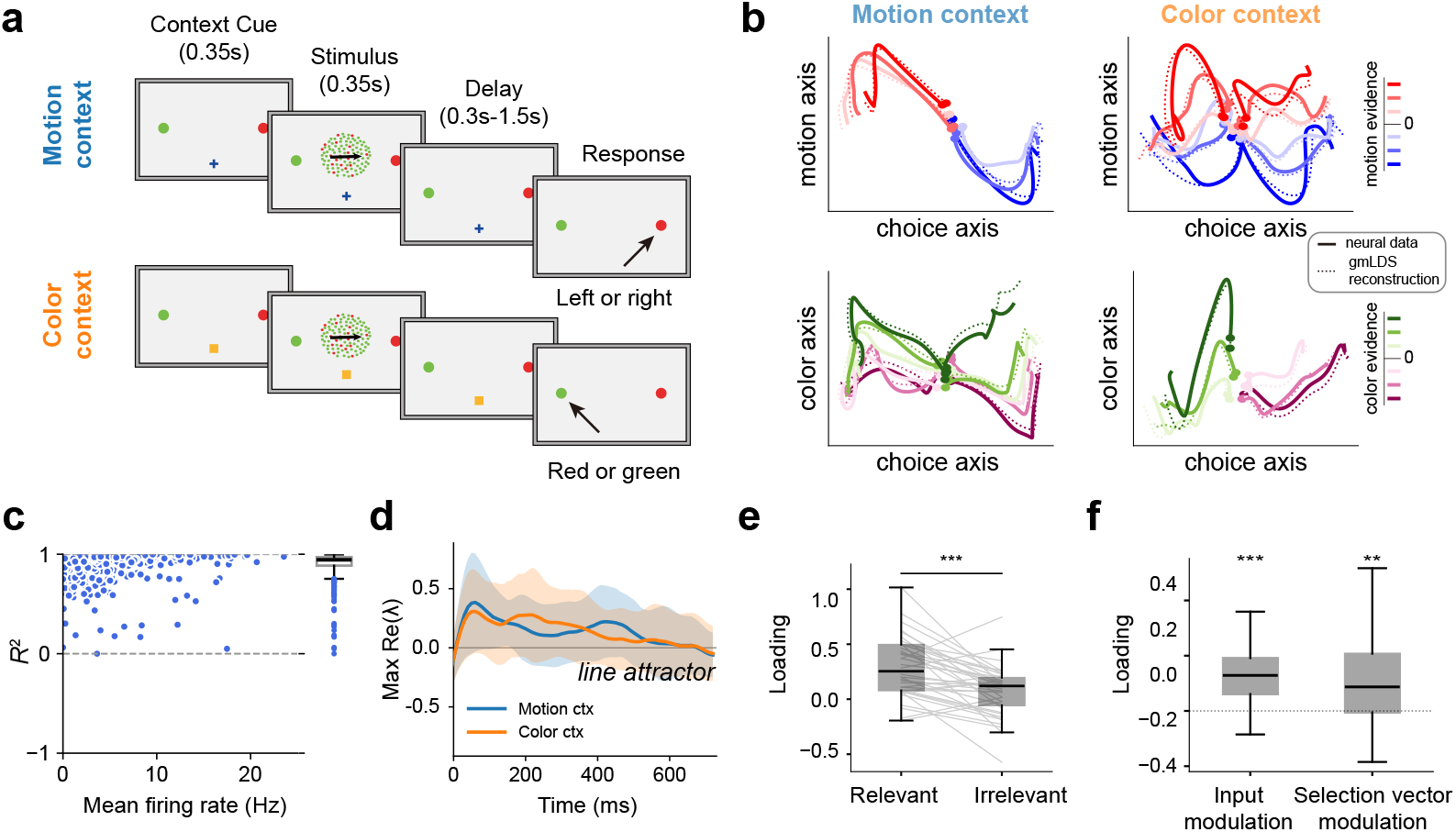
gmLDS reveals context-dependent computational mechanisms in primate pre-frontal cortex. **a**, Task paradigm for CDM task. **b**, Low-dimensional population trajectories projected onto task-relevant subspaces identified using targeted dimensionality reduction (TDR). gmLDS accurately captures the geometry of neural population activity across task contexts. **c**, Model fit quality quantified by the coefficient of determination (*R*^2^) across neurons. Each point corresponds to a single neuron; mean *R*^2^ = 0.901 (s.d. = 0.129). **d**, Time course of the largest real eigenvalue of the inferred effective connectivity matrix, max Re(*λ*), aligned to stimulus onset. Shaded regions denote mean ± s.e.m. across conditions. In both motion and color contexts, the dominant eigenvalue remains close to zero during the stimulus period, consistent with line-attractor–like dynamics. **e**, Projection of input representation onto the selection vector in relevant and irrelevant contexts. The projection is significantly larger in the relevant context, indicating selective accumulation of context-relevant information. (*t*-test, *p <* 0.001, *n* = 40). **f**, Input modulation and selection-vector modulation inferred by gmLDS. Bars denote mean ± s.e.m. across neurons. Both modulation terms are significantly greater than zero (*t*-test, \*\**p <* 0.01, \*\*\**p <* 0.001).

**Figure 5:**
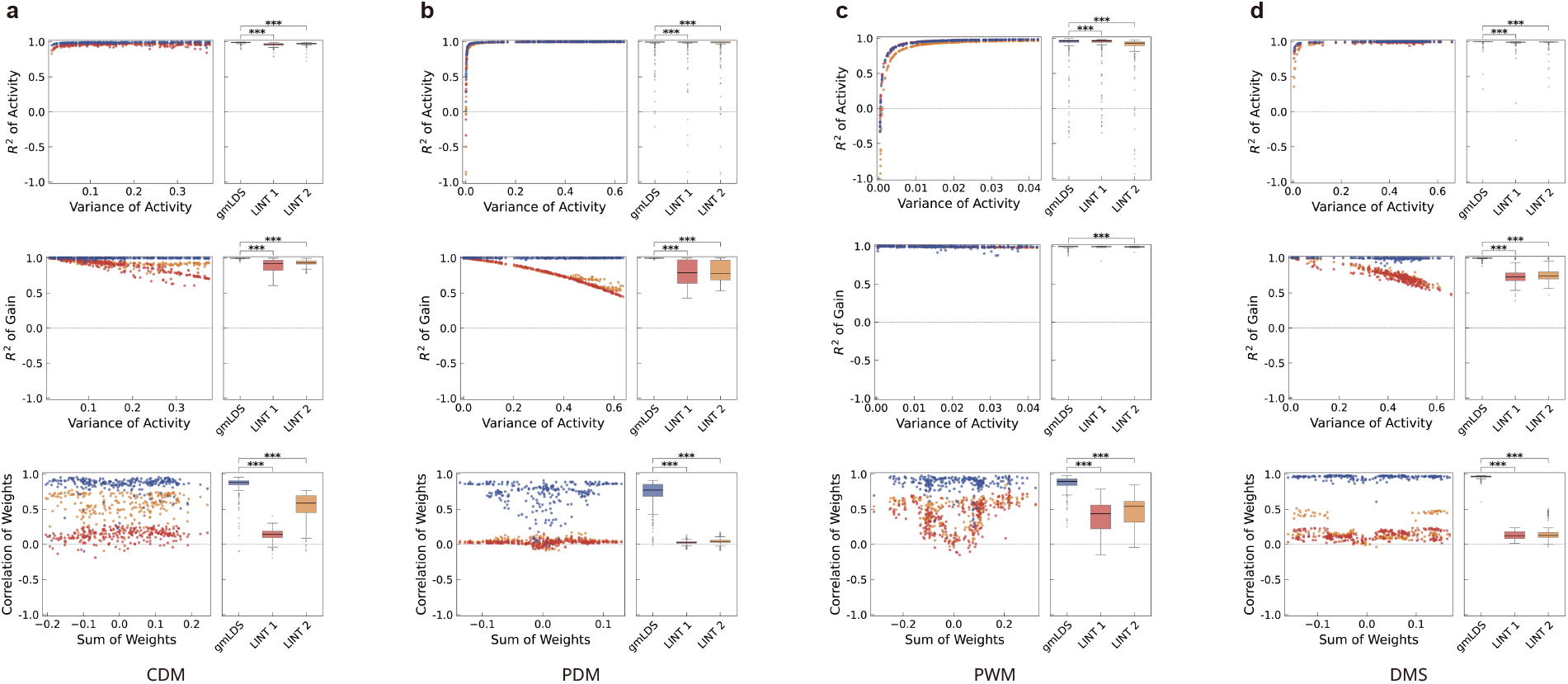
Neuron-wise performance versus activity variance and input-weight magnitude across tasks. For each task, the left panels show per-neuron scatter plots and the right panels summarize across-neuron distributions (boxplots). Top: activity *R*^2^ vs. activity variance. Middle: gain *R*^2^ vs. activity variance. Bottom: correlation of recovered vs. ground-truth input weights vs. sum of input weights. Brackets denote paired Wilcoxon signed-rank tests vs. gmLDS (* *p <* 0.05, ** *p <* 0.01, *** *p <* 0.001). **a** CDM, **b** PDM, **c** PWM, **d** DMS.

**Figure 6:**
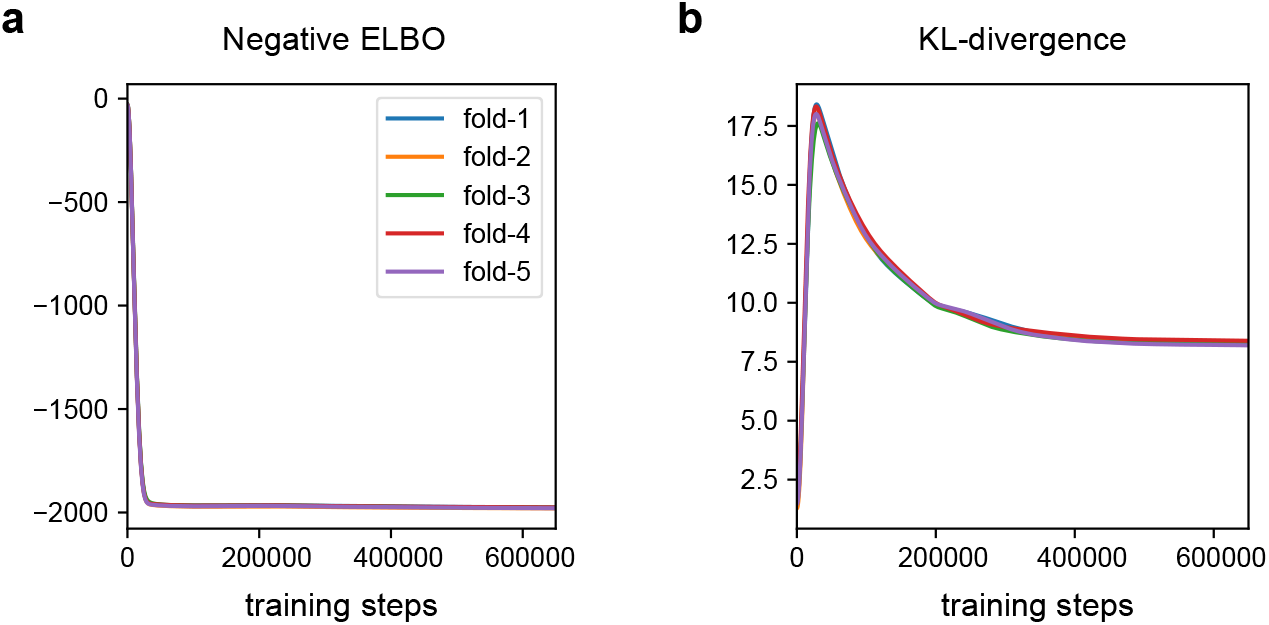
Loss throughout training process in fitting monkey data.

**Figure 7:**
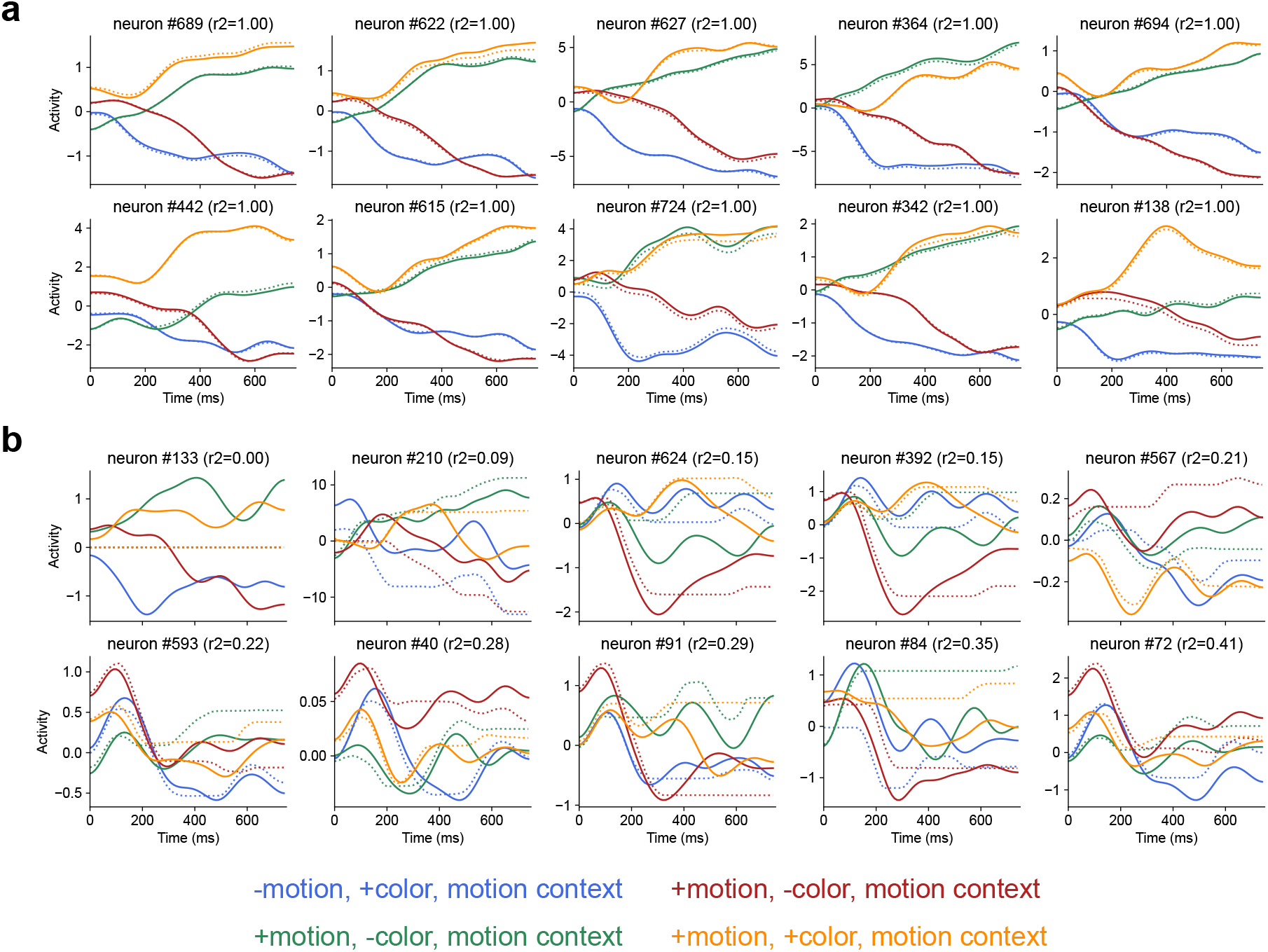
Reconstruction of single-neuron activity. **a**, The 10 neurons with the highest reconstruction accuracy (*r*^2^). **b**, The 10 neurons with the lowest reconstruction accuracy. For each neuron, activity in four trials with the strongest stimulus coherence is displayed. Solid lines denote ground-truth activity, and dotted lines denote model reconstructions. Colors indicate the four stimulus conditions, as defined below.

**Figure 8:**
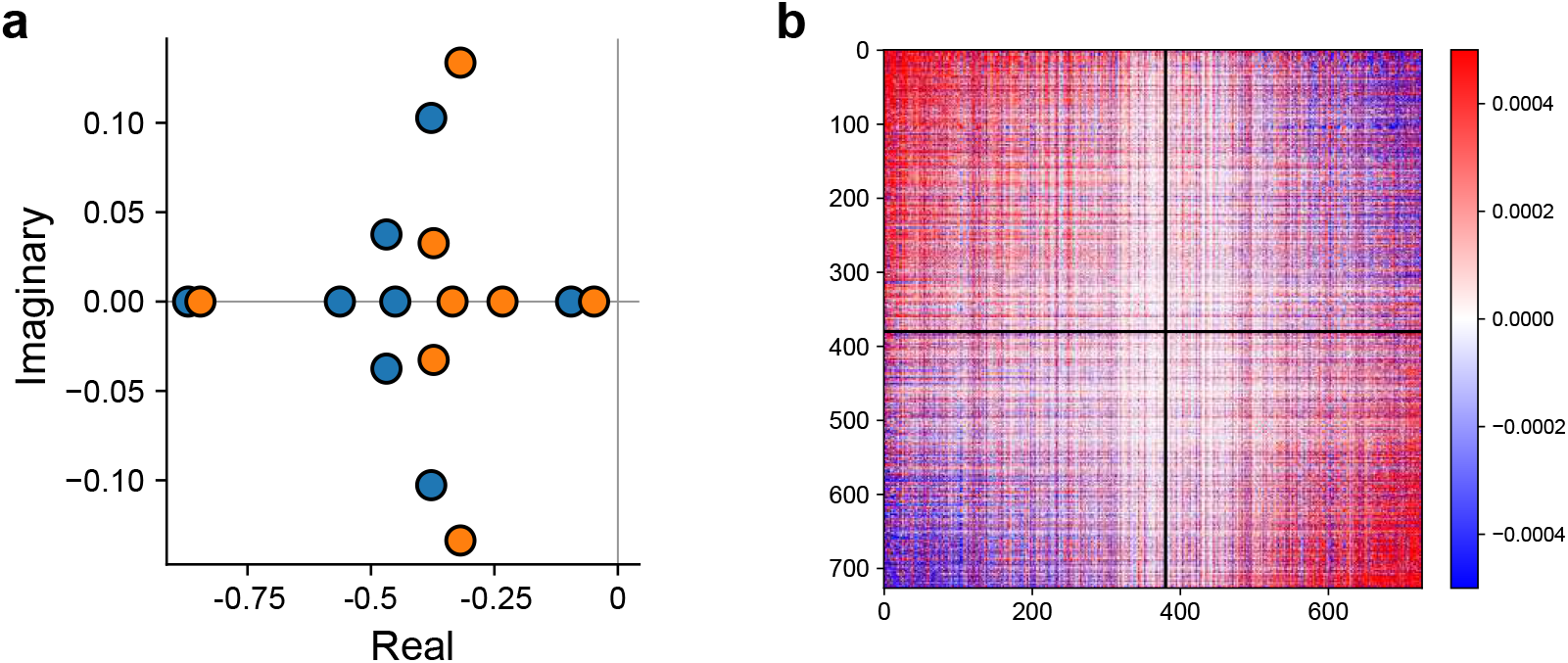
**a**, Eigenvalue spectrum of the linearized RNN dynamics in the complex plane. Blue and orange markers correspond to context 1 and context 2, respectively. **b**, Connectivity matrix inferred by *mn*^*T*^. Neurons are sorted by their preference of the motion stimulus given by TDR.

We focused on population-level dynamics during the stimulus period. Neural responses were pre-processed to obtain smooth, trial-averaged firing-rate trajectories for each task condition, and projected onto a low-dimensional subspace capturing the dominant population activity patterns before reconstruction in the original neural space (see Appendix 9.1 for details).

The model received externally specified sensory and contextual input channels corresponding to task structure, and was trained to reconstruct neural activity during the stimulus period. This setup allowed gmLDS to infer time-varying gain modulation and low-rank effective connectivity directly from neural activity, enabling us to characterize the underlying dynamical structure and its modulation by task context.

We traind and evaluated gmLDS on this dataset using a five-fold cross-validation (5-fold CV) co-smoothing scheme (Pei et al. 2021). For each fold, neural activity from four folds was provided to the posterior inference model to infer the posterior correction term *w*_*t*_. Using this inferred posterior, the log-likelihood–based reconstruction loss was computed and optimized over all neurons, while the remaining fold was held out for evaluation. Model was trained with mini-batches of size 216 for 650K steps, corresponding to three trials per condition across all task conditions. The learning rate was 0.0005. To prevent the learned dynamics from being dominated by the posterior correction term and to encourage interpretable latent dynamics, we introduced a weighting factor *β* on the KL-divergence term in the ELBO objective, setting *β* = 2. This choice places slightly greater emphasis on inferring structured latent vector fields at the cost of marginally reduced reconstruction accuracy, a trade-off that has been shown to promote interpretable latent representations in variational models (Higgins et al. 2017).

We next assessed the fitting performance of gmLDS on this dataset to evaluate how accurately the model inferred neural responses from held-out data. At the single-neuron level, gmLDS achieved consistently high reconstruction accuracy across neurons (Fig. 4c), demonstrating that the model captured heterogeneous response profiles despite substantial variability in firing patterns. At the population level, we examined the geometry of inferred neural trajectories by projecting both recorded and gmLDS-inferred activity onto task-relevant axes identified using targeted dimensionality reduction (TDR) (Mante et al. 2013) (Fig. 4b). The close correspondence between inferred and ground-truth population trajectories indicates that gmLDS successfully recovered the low-dimensional neural dynamics underlying context-dependent computation in the CDM task.

To characterize the dynamical regime underlying the inferred population activity, we examined the spectrum of the Jacobian matrix recovered by gmLDS. The largest real eigenvalue remained close to zero throughout the stimulus period in both motion and color contexts (Fig. 4d). We further examined the full eigenvalue spectrum at stimulus onset to characterize the dynamical structure in greater detail (Supp. Fig. 8a). Aside from a single eigenvalue with a real part close to zero, all remaining eigenvalues had strictly negative real parts. Moreover, a subset of eigenvalues exhibited non-zero imaginary components, suggesting the presence of rotational components in the population dynamics. Together, these spectral properties indicate that neural activity in each context during the CDM task is organized around a low-dimensional line-attractor structure, consistent with theoretical explanation for context-dependent computation.

Building on this dynamical characterization, we defined the gmLDS *selection vector* as the right eigenvector associated with the eigenvalue having the largest real part of the Jacobian matrix evaluated at stimulus onset. To quantify the direction of sensory input in population space, we computed the input representation direction as 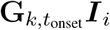, where 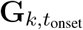 denotes the inferred gain at stimulus onset for trial *k*, and ***I***_*i*_, *i* = 1, 2 is the input embedding for each modality.

We next quantified the alignment between the input representation and the selection vector across the two task contexts. For each context, we analyzed trials in which both sensory modalities exhibited large absolute evidence strength, yielding four stimulus conditions per context. Results were aggregated across five cross-validation folds, resulting in 40 samples per context (2 modalities × 4 conditions × 5 folds). We found that this alignment was significantly stronger in the task-relevant context than in the irrelevant context, indicating that sensory evidence is preferentially projected onto the selection vector when it is behaviorally relevant, thereby supporting more effective evidence accumulation (Fig. 4e).

Finally, we asked whether contextual modulation in the CDM task manifests through input modulation, selection-vector modulation, or both. To address this question, we quantified these two forms of modulation captured by gmLDS, as defined in Eq. 16. We found that both modulation terms were significantly greater than zero (one-sample, one-sided *t*-tests; Fig. 4f). These results indicate that task context influences context-dependent computation through both modulation of effective sensory inputs and altering of selection vectors, providing evidence that both mechanisms coexist in the macaque CDM task.

## Discussion

A central contribution of gmLDS is to enable direct inference of local linearized dynamics from population recordings alone, without imposing strong parametric assumptions on the underlying neural model. This effectively extends the classical “opening the black box” method (Sussillo et al. 2013), which rely on computing Jacobian matrices of fully specified RNNs, thereby requiring access to the trained model and its activation function. In this sense, gmLDS represents a principled step from post-hoc inspection to data-driven discovery of computational mechanism from neural recordings.

Beyond its methodological contribution, gmLDS also helps reconcile seemingly divergent conclusions in the literature regarding the mechanisms underlying context-dependent computation. In the original work (Mante et al. 2013), context-dependent behavior was explained through modulation of the selection vector, based on analyses of a task-driven RNN. Subsequent studies adopting low-rank modeling approaches, however, demonstrated that neural activitiy in this task could be captured by rank-1 networks, implicitly favoring an input-modulation interpretation (Valente et al. 2022). Later work (Soldado-Magraner et al. 2024) further showed that either context-dependent changes in recurrent dynamics or in input representations could explain prefrontal neural activity equally well, leading to the conclusion that these mechanisms are non-identifiable from neural data alone.

By explicitly modeling gain-modulated connectivity and the resulting linearized dynamics, gmLDS provides a unified, data-driven framework that reframes this identifiability problem. In gmLDS, input modulation and transient dynamical modulation arise from a shared gain factor, rather than being treated as mutually exclusive mechanisms. This formulation changes the identifiability structure of the model and allows both modulations to be quantitatively assessed directly from neural recordings. Applying this framework, we find evidence for the coexistence of these two selection mechanisms, offering a principled resolution to a long-standing debate on context-dependent computation.

gmLDS is also complementary to a broad class of latent dynamical inference methods, including LFADS, rSLDS, FINDER (Pandarinath et al. 2018; Linderman et al. 2017; Kim et al. 2025). These approaches excel at capturing latent trajectories and predicting neural activity, but typically operate at a latent or algorithmic level, without explicitly linking the inferred dynamics to neuron-level interactions. In contrast, gmLDS provides an implementational-level description in which population dynamics arise from gain-modulated interactions between neurons, enabling direct interpretation of how connectivity structure gives rise to observed dynamics. Moreover, the gain-modulated LDS formulation has a close mathematical correspondence to the local linearization of low-rank RNNs, providing a principled bridge between data-driven dynamical inference and theoretical analyses of recurrent networks.

Several extensions could further broaden the applicability of gmLDS, including inferring external inputs rather than specifying them a priori and incorporating observation models tailored to spike-train data. Applying the framework to more complex cognitive tasks with higher-dimensional and richer neural dynamics, as well as extending it to integrate neural recordings across days or individuals, remains challenging and represents an important direction for future work. Progress along these lines may further strengthen gmLDS as a bridge between data-driven dynamical analyses and mechanistic theories of neural computation.

## Appendix

### Synthetic neural data on multiple cognitive tasks

We adopt the four cognitive tasks: perceptual decision-making (PDM) (Gold et al. 2007), contextual decision-making (CDM) (Mante et al. 2013), parametric working memory (PWM) (Romo et al. 1999), and delayed match-to-sample (DMS) (Miyashita 1988). During simulation, we adopt the same setting as Dubreuil et al. 2022.

#### Perceptual decision-making (PDM)

The task has four consecutive phases—fixation (100 ms), stimulus (800 ms), delay (100 ms), response (20 ms). A coherence *c* is drawn uniformly from {−4, −2, −1, 1, 2, 4}. Inputs are one-dimensional; in the stimulus phase we set stimulus *u*_*t*_ = *s*_stim_ · *c* + *s*_noise_ · *ξ*_*t*_ with scale *s*_stim_ = 0.1, *s*_noise_ = 0.1 and i.i.d. Gaussian noise *ξ*_*t*_. The target *y*_*t*_ equals sign(*c*) in the response phase and is zero otherwise. The mask *H*_*t*_ is 1 in the response window only when *c*≠ 0, so zero-coherence trials are excluded from the loss.

#### Contextual decision-making (CDM)

The task has five consecutive phases—fixation (100 ms), context (350 ms), stimulus (800 ms), delay (100 ms), response (20 ms). We sample a context cue *c*_ctx_ ∈ {0, 1} and coherences *c*_*A*_, *c*_*B*_ ∈ {−4, −2, −1, 1, 2, 4}. Inputs are four-dimensional: two evidence channels and two context channels. In the stimulus phase we set *u*_0_ = *s*_stim_ · *c*_*A*_ + *s*_noise_ · *ξ*_*A*_(*t*) and *u*_1_ = *s*_stim_ · *c*_*B*_ + *s*_noise_ · *ξ*_*B*_(*t*) with *s*_stim_ = 0.1, *s*_noise_ = 0.1 and i.i.d. Gaussian *ξ*_*A*_, *ξ*_*B*_; outside this phase the evidence channels are zero. From context onset to trial end one context channel is *s*_ctx_ = 0.1, the other 0 (one-hot). The target *y*_*t*_ equals sign(*c*_*A*_) in the response phase when *c*_ctx_ = 0 and sign(*c*_*B*_) when *c*_ctx_ = 1, and is zero otherwise. The mask *H*_*t*_ is 1 in the response window only when the cued coherence is non-zero.

#### Parametric working memory (PWM)

The task has five phases—fixation (100 ms), first stimulus (100 ms), ISI uniformly sampled in [500, 1000] ms, second stimulus (100 ms), response (100 ms). We draw a frequency difference Δ*f* ∈ {−24, −16, −8, 8, 16, 24} and choose *f*_base_ so that *f*_1_ = *f*_base_ and *f*_2_ = *f*_base_ + Δ*f* lie in [10, 34], then set *u*_1_ = (*f*_1_ − 22)*/*24 and *u*_2_ = (*f*_2_ −22)*/*24. Inputs are one-dimensional; in the first stimulus phase we set *u*_*t*_ = *u*_1_ + *s*_noise_ · *ξ*_*t*_, in the second stimulus phase *u*_*t*_ = *u*_2_ + *s*_noise_ · *ξ*_*t*_, with *s*_noise_ = 0.01 and i.i.d. Gaussian *ξ*_*t*_; elsewhere *u*_*t*_ is zero plus noise. The target *y*_*t*_ equals *u*_1_ − *u*_2_ in the response phase and is zero otherwise. The mask *H*_*t*_ is 1 in the response window except when *u*_1_ = *u*_2_, in which case that trial is excluded from the loss.

#### Delayed match-to-sample (DMS)

The task has five phases—fixation (100 ms), first stimulus (500 ms), delay uniformly sampled in [500, 3000] ms, second stimulus (500 ms), response (1000 ms). Inputs are two-dimensional; in each stimulus phase we draw *s* ∈ {*A, B*} and set (*u*_0_, *u*_1_) = (1, 0) for A or (0, 1) for B. At all times we add *s*_noise_ · *ξ*_*t*_ with *s*_noise_ = 0.03 and i.i.d. Gaussian *ξ*_*t*_ to both channels, and outside stimulus phases the two channels are zero plus noise. The target *y*_*t*_ equals +1 when the two cues match and −1 when they do not in the response phase and is zero otherwise. The mask *H*_*t*_ is 1 in the response phase and 0 elsewhere.

#### Training ground-truth RNNs

For each cognitive task we train a family of vanilla recurrent neural networks with *N* = 256 units. The networks receive the task inputs *u*_*t*_ defined above and produce a scalar output at each time step. The hidden state *h*_*t*_ ∈ ℝ^*N*^ follows a discrete-time dynamics

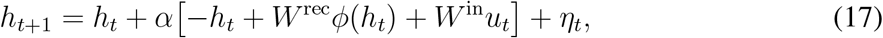

with integration constant *α* = 0.2, dynamic noise 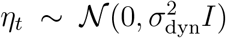 with *σ*_dyn_ = 10^−2^, and activation function *ϕ*. The readout is linear in the nonlinearity, *ŷ*_*t*_ = *W* ^out^*ϕ*(*h*_*t*_)*/N*.

We consider both tanh(·) and softplus(·) −1 as activation functions. Given target *y*_*t*_ and mask *H*_*t*_ defined above, networks are trained to minimize a masked mean-squared error

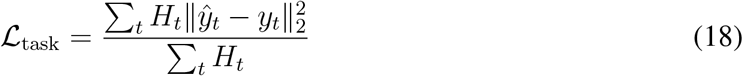

plus an *ℓ*_2_ penalty on the recurrent weights, 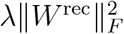. The regularization strength *λ* is task-dependent (on the order of 10^−4^) and chosen so that the trained connectivity implements a low-dimensional dynamics (effective rank about 1–4) while preserving high decision accuracy.

For all tasks we train the RNNs directly from the task generators using the Adam optimizer, with mini-batches of size 512, learning rate 10^−3^, gradient clipping (between 5 × 10^−3^ and 10^−2^ depending on the task), and early stopping on a held-out validation set, for at most 10^4^ epochs. For each task and hyperparameter setting we repeat training 8 times with different random seeds and use all trained networks for subsequent analysis.

### LINT fitted to synthetic neural data

To analyze the dynamics of the task-optimized RNNs, we use LINT (Low-rank Inference from Neural Trajectories) as a low-rank recurrent network that directly fits neural activity. LINT shares the same dynamical form as the vanilla RNN described above, including the integration constant *α* = 0.2, but is trained in “activity” mode to predict hidden states (or their nonlinear activations) of a ground-truth network, rather than task outputs. The effective recurrent connectivity of LINT is constrained to be low-rank, with rank *r* ∈ {1, 2, 4} depending on the configuration.

We fit LINT to synthetic neural data generated by the task-trained RNNs on the four cognitive tasks. For each task we consider three LINT variants distinguished by their activation function: **LINT 1** uses softplus(·) −1, **LINT 2** uses 2 retanh(·) −1, and **LINT 3** uses tanh(·). The rectified tanh is defined as

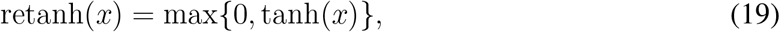

and the variant 2 retanh(*x*) −1 keeps the output approximately in [−1, 1] while enforcing rectification.

LINT models are trained directly on RNN-generated neural trajectories using the Adam optimizer, with mini-batch size 512, learning rate 10^−3^, gradient clipping in the range 10^−2^, and early stopping on a held-out validation set. For each task and hyperparameter configuration we train eight independent LINT instances with different random seeds and include all of them in the downstream analysis.

We evaluate how well LINT recovers the synthetic neural dynamics and associated computational mechanisms. As summarized in Supp. Table. 1, we report coefficients of determination (*R*^2^) between predicted and ground-truth neural activity and gain trajectories, together with Pearson correlation coefficients between inferred and ground-truth low-rank connectivity vectors.

### gmLDS implementation details

#### Neural modules

The gmLDS implementation uses three neural components: an encoding network, a correction network, and a gain network. The encoding network first concatenates neural activity and stimulus input at each time step, then passes this combined signal through a small two-layer feedforward network with 64 units in each hidden layer, and finally feeds the compressed sequence into a two-layer unidirectional LSTM with 64 hidden units. The LSTM is run backward in time and its outputs are flipped, so that at each time point we obtain a context vector that summarizes future neural and stimulus increments.

The correction network takes as input the context vector together with the current latent state and outputs the parameters of a Gaussian distribution over the process noise at that time step. It consists of a shared fully connected layer with 64 hidden units and a sigmoid nonlinearity, followed by three shallow heads that map this shared representation to the mean, the diagonal log-variance, and an optional low-rank factor of the noise covariance. The dimension of the noise vector is set equal to the latent dimension by default. When the low-rank factor is disabled, the covariance reduces to a purely diagonal form.

The gain network maps the latent state and the integrated stimulus into a time-varying gain profile over neurons. It concatenates these two inputs and applies layer normalization followed by a three-layer multilayer perceptron with two hidden layers of width 512 and LeakyReLU activations, together with a residual skip connection from input to output scaled by a learnable scalar. A final softplus nonlinearity ensures that the predicted gains are non-negative, and these gains are used as a diagonal modulation of a static low-rank connectivity and input matrix to form the time-varying operators *A*_*t*_, *B*_*t*_, *C*_*t*_, *D*_*t*_.

#### Noise modeling with LRPD

We parameterize both the observation noise covariance *Q* and the process noise covariance *R* using a low-rank-plus-diagonal (LRPD) family. This formulation guarantees positive definiteness, allows efficient computation of inverses and log-determinants via the Woodbury identity and matrix determinant lemma, and supports a straightforward reparameterization trick for sampling.

For a covariance matrix of dimension *d* and chosen rank *ρ* (typically *ρ* = 1 in our experiments), we write

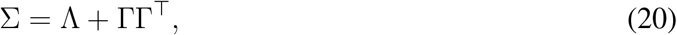

where Λ = diag(*λ*_1_, …, *λ*_*d*_) is a diagonal matrix with strictly positive entries and Γ ∈ ℝ^*d*×*ρ*^ is a low-rank factor with *ρ* ≪ *d*. In practice we parameterize Λ via unconstrained log-variances, *λ*_*i*_ = exp(*ℓ*_*i*_), and learn Γ directly.

Using the Woodbury matrix identity(Woodbury 1950), the inverse of Σ can be written as

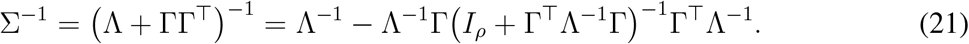

When *ρ* = 1, Γ reduces to a single column vector *γ* ∈ ℝ^*d*^ and the inner term becomes a scalar,

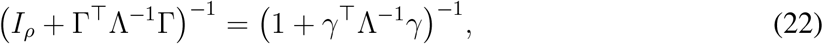

so that only one scalar inversion is required.

Similarly, by the matrix determinant lemma we have

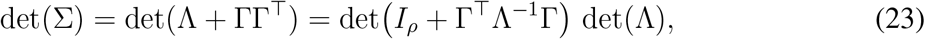

and for *ρ* = 1 this simplifies to

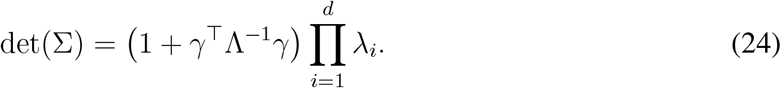

For sampling and the reparameterization trick, we use the equivalent factorization

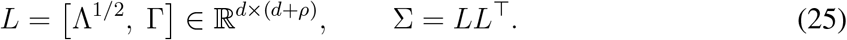

Given standard normal noise *ε* ∼ 𝒩 (0, *I*_*d*+*ρ*_), we draw

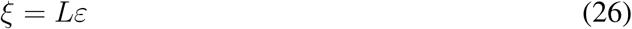

so that *ξ* ∼ 𝒩 (0, Σ). Within gmLDS we maintain separate LRPD parameterizations for *Q* and *R*, allowing both anisotropic marginal variances and low-rank correlations in observation and latent noise, while very low ranks (most often *ρ* = 1) are sufficient to capture the dominant correlated variability with modest computational cost.

#### Generation algorithm

Sampling from the gmLDS generative model proceeds by forward simulation in the differential domain. Given an input sequence 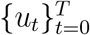, we first compute the integrated input *ν*_*t*_ and increments Δ*ν*_*t*_ by leaky integration with time constant *τ*. We initialize a baseline activity *x*_0_ and a latent state *z*_0_ (e.g., both zeros). For each time step *t* = 1, …, *T* we then:

1. Update the accumulated latent state 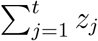 and evaluate the gain *G*_*t*_ via the gain module.
2. Assemble *A*_*t*_, *B*_*t*_, *C*_*t*_, *D*_*t*_ from *G*_*t*_, *m, n, I*.
3. Sample process noise *w*_*t*_ ∼ 𝒩 (0, *R*) and observation noise *ϵ*_*t*_ ∼ 𝒩(0, *Q*) using the LRPD samplers.
4. Update the latent increments

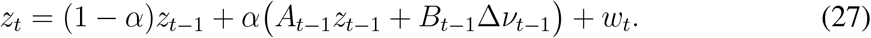
5. Compute the predicted activity increment

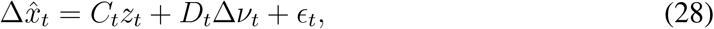

and update 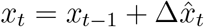.

The result is a synthetic neural trajectory 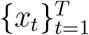 together with latent increments {*z*_*t*_} and gain profiles {*G*_*t*_} consistent with the gmLDS prior.

#### Inference algorithm

Posterior inference in gmLDS follows a similar forward-simulation logic, but with the process noise at each time step drawn from a learned posterior instead of the prior. Given observed activity and stimulus increments, we proceed as follows:

1. Run the encoding network on the full sequences of neural activity and stimulus input to obtain a sequence of context vectors. Concretely, the compressed (*x*_*t*_, *u*_*t*_) sequence is passed through the backward LSTM and its outputs are reversed to produce

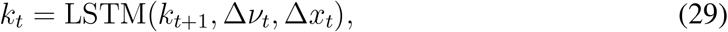

where the backward LSTM processes the sequence from *t* to *T*.
2. Initialize the latent state *z*_0_ using the learned initialization network, which maps an auxiliary noise sample and the initial input into the latent space.
3. For each time step *t*, feed (*z*_*t*−1_, *k*_*t*_) into the correction network to obtain the mean and LRPD covariance parameters of a Gaussian posterior over the process noise,

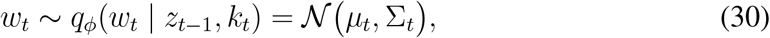

draw a sample via reparameterization,

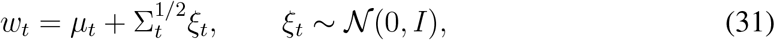

and update the latent state using the same linear dynamical rule as in the generative model,

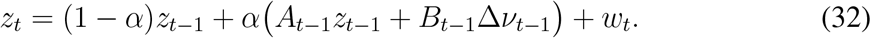
4. For each time step, pass *z*_*t*_ through the generative part of the model to obtain a predicted increment 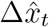 and its reconstruction log-likelihood under *p*_*θ*_(Δ*x*_*t*_|*z*_*t*_, Δ*ν*_*t*_), and compute the KL regularization term

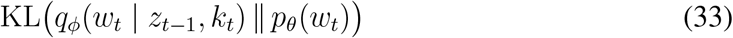

between the learned posterior over the process noise and its prior.
5. Sum reconstruction and KL terms over time to obtain the variational evidence lower bound, and optimize all model parameters by backpropagation through time using one Monte Carlo sample of the process noise per sequence.

The learned model provides time-varying gains and static low-rank connectivity factors that we compare against the ground-truth low-rank RNNs in the main text.

### gmLDS fitted to synthetic neural data

In addition to LINT, we also fit gmLDS models directly to the synthetic neural activity generated by the task-trained RNNs on all four cognitive tasks. For each ground-truth RNN, we train gmLDS variants with latent dimensions 1, 2, and 4 (chosen according to the task settings), setting *α* = 0.2, using the Adam optimizer with mini-batch size 512, learning rate 10^−3^, gradient clipping of order 10^−2^, and early stopping on a validation set. The same compact metrics as for LINT—in particular the *R*^2^ between gmLDS-predicted and ground-truth neural activity and gain trajectories, and Pearson correlation between inferred and ground-truth connectivity vectors—are summarized in Supp. Table. 1.

Supp. Table. 1 summarizes the performance of gmLDS and LINT across the four cognitive tasks (CDM, PDM, PWM, DMS) and two ground-truth activation functions (tanh(·) and softplus(·)−1). All models achieve high *R*^2^ of activity fitting, indicating that neural population dynamics can be accurately reconstructed by both approaches. However, LINT fails to recover the ground-truth neural gain when its activation function is mismatched to the ground-truth RNN: under tanh(·) ground truth, LINT 1 (softplus(·) −1) and LINT 2 (2 retanh(·) −1) yield low *R*^2^ of gain inferring on CDM, PDM, and DMS, whereas gmLDS consistently infers the gain well across all tasks. Similarly, under softplus(·) −1 ground truth, LINT 2 (2 retanh(·) −1) and LINT 3 (tanh(·)) perform poorly on gain inference, while gmLDS maintains high gain *R*^2^. On connectivity, gmLDS substantially outperforms LINT, with higher Pearson correlation between inferred and ground-truth recurrent weights. One notable exception is the PWM task: on this task, all models—including LINT with mismatched nonlinearity—achieve high *R*^2^ of gain inferring. We suspect that in PWM the overall variance of neural activity in the ground-truth network is relatively low (Supp. Fig. 5c), leading to low variance in the gain profile as well; this may make the gain easier to infer regardless of model structure, and thus attenuate the advantage of gmLDS on this particular task.

**Table 1:** Performance comparison across tasks and ground-truth activation functions.

| Ground-truth activation | Model | CDM | PDM | PWM | DMS |
| --- | --- | --- | --- | --- | --- |
| tanh(·) | *R<sup>2</sup> of activity fitting* | | | | |
|  | gmLDS | **.9911±.0012** | **.9981±.0003** | **.9694±.0015** | **.9976±.0005** |
| | LINT 1 | .9628±.0011 | .9972±.0003 | .9672±.0015 | .9898±.0021 |
| | LINT 2 | .9682±.0109 | .9927±.0079 | .9394±.0036 | .9932±.0017 |
|  | *R<sup>2</sup> of gain inferring* |  |  |  |  |
| | gmLDS | **.9953±.0008** | **.9977±.0003** | .9909±.0020 | **.9878±.0029** |
| | LINT 1 | .8874±.0068 | .7992±.0222 | **.9922±.0005** | .7499±.0336 |
| | LINT 2 | .9266±.0060 | .8130±.0228 | .9911±.0006 | .7695±.0373 |
|  | *Correlation of connectivity* |  |  |  |  |
|  | gmLDS | **.8302±.0208** | **.7008±.0476** | **.8747±.0210** | **.9454±.0138** |
| | LINT 1 | .1639±.0261 | .0243±.0036 | .3555±.0380 | .1781±.0545 |
| | LINT 2 | .4799±.0909 | .0377±.0037 | .4758±.0598 | .2750±.1044 |
| softplus(·) - 1 | *R<sup>2</sup> of activity fitting* | | | | |
|  | gmLDS | **.9988±.0002** | **.9996±.0001** | **.9943±.0005** | **.9995±.0002** |
| | LINT 2 | .3235±.0218 | .3625±.0499 | .8785±.0122 | .3479±.0315 |
| | LINT 3 | .3356±.0232 | .4548±.0426 | .8991±.0114 | .3538±.0335 |
|  | *R<sup>2</sup> of gain inferring* |  |  |  |  |
|  | gmLDS | **.9844±.0059** | **.9890±.0026** | **.9959±.0008** | **.9931±.0041** |
| | LINT 2 | .6987±.0174 | .7448±.0239 | .9432±.0025 | .6801±.0304 |
| | LINT 3 | .7897±.0158 | .8583±.0217 | .9706±.0023 | .8060±.0233 |
|  | *Correlation of connectivity* |  |  |  |  |
|  | gmLDS | **.6924±.0330** | **.7596±.0238** | **.8461±.0503** | **.7446±.0363** |
| | LINT 2 | .0900±.0310 | .0399±.0430 | .1682±.0144 | .0676±.0393 |
| | LINT 3 | .1250±.0264 | -0.0016±.0273 | .2056±.0233 | .1052±.0448 |

### Linearized dynamical system analysis of RNNs around fixed point

In this appendix, we briefly summarize the linearized dynamical system analysis used to interpret RNNs (Mante et al. 2013). Throughout, we denote neural firing rates by ***x***, with ***x*** = *ϕ*(***h***), where ***h*** is the pre-activation and *ϕ*(·) is an elementwise nonlinearity, following the notation in the main text.

An RNN can be written as a nonlinear dynamical system

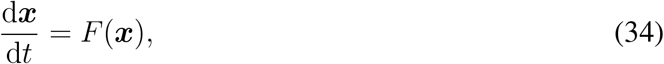

where ***x*** ∈ ℝ^*N*^ and *F* : ℝ^*N*^ → ℝ^*N*^. Consider a fixed point ***x***^∗^ satisfying *F* (***x***^∗^) = **0**. Writing *δ****x*** = ***x*** − ***x***^∗^, a first-order Taylor expansion yields

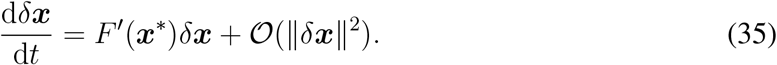

Neglecting higher-order terms for sufficiently small perturbations leads to the local linear system

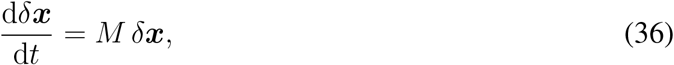

where *M* = *F* ^′^(***x***^∗^) ∈ ℝ^*N* ×*N*^ is the Jacobian evaluated at the fixed point.

Assuming *M* is diagonalizable, its eigendecomposition can be written as

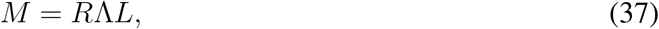

where Λ = diag(*λ*_1_, …, *λ*_*N*_) contains the (generally complex) eigenvalues, and *R* and *L* are the corresponding right and left eigenvector matrices, satisfying *R*^−1^ = *L*. Introducing coordinates ***y*** = *Lδ****x*** decouples the dynamics:

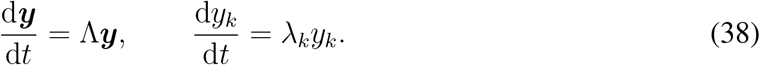

Each mode admits the solution 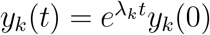, and transforming back gives

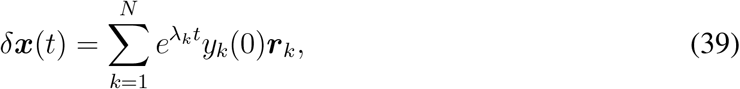

where ***r***_*k*_ is the right eigenvector associated with *λ*_*k*_.

For real eigenvalues *λ*_*k*_ ∈ ℝ, perturbations grow or decay exponentially along ***r***_*k*_. For complex-conjugate eigenvalue pairs *λ*_*k*_ = *σ*_*k*_ ± *iω*_*k*_, the dynamics correspond to oscillatory modes with envelope 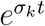. The spectrum of *M* therefore determines the local behavior near ***x***^∗^, including attracting, repelling, saddle, and oscillatory regimes.

### Details for synthetic CDM task

#### Task settings

##### Click-based CDM task

We simulated a click-based context-dependent decision-making task recently investigated by Pagan et al. 2024. Each trial consisted of a fixation period (200 ms), a stimulus period (800 ms), and a decision period (20 ms). During the stimulus period, sensory evidence was delivered as stochastic pulse trains. At each time step, the total number of pulses was drawn from a Poisson distribution with mean 0.8. Each pulse was independently assigned a spatial attribute (left or right) and a feature attribute (low or high frequency), with trial-specific probabilities *p*_*k*_ and 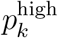, respectively. These probabilities were independently sampled from the set {39*/*40, 35*/*40, 25*/*40, 15*/*40, 5*/*40, 1*/*40}.

The task contains four input channels *u*_1_(*t*), *u*_2_(*t*), 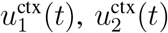. The first sensory input *u*_1_(*t*) encoded spatial evidence as the scaled difference between rightward and leftward pulses, and the second sensory input *u*_2_(*t*) encoded frequency evidence as the scaled difference between high- and low-frequency pulses. The context inputs 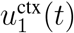 and 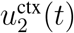 indicate whether the current context is context-1 or context-2. In context-1, 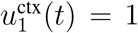 and 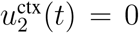 throughout the entire trials. Similarly, in context-2, 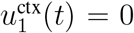 and 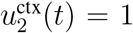 throughout the entire trials. Task context was randomly assigned on each trial and was represented by constant contextual inputs that indicated which sensory modality was behaviorally relevant. In the spatial context, the target output depended only on the sign of the spatial stimulus strength 2*p*_*k*_ − 1, whereas in the frequency context it depended only on the sign of the frequency stimulus strength 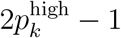.

#### Training of ground-truth vanilla RNNs

To evaluate gmLDS in a controlling setting, we generate synthetic data using vanilla RNNs trained on the click-based CDM task following established protocols. Specifically, we trained vanilla RNNs with *N* = 128 neurons, whose dynamics are governed by

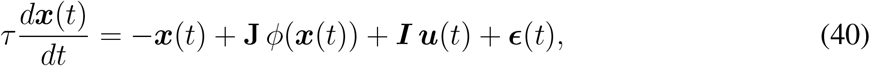

where ***x***(*t*) denotes the neural state, **J** the recurrent connectivity, *ϕ* = tanh a nonlinear activation function, ***u***(*t*) the external input, and ***ϵ***(*t*) is a time-varying noise term. The network received two sensory input channels corresponding to two stimulus modalities, together with two context input channels indicating which modality was relevant on each trial (Fig. 3a).

The network’s output was read out from the population activity ***x***(*t*) through a linear mapping:

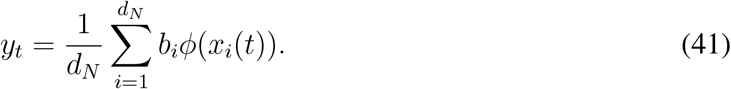

All recurrent connections, input weights, and readout weights were optimized during training. The loss function took the form:

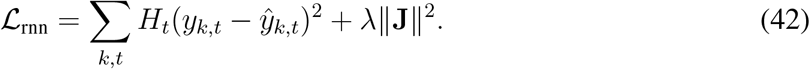

where *y*_*k,t*_ is the target output and *H*_*t*_ is a {0, 1} mask being 1 in the decision period and 0 otherwise. The connectivity matrix *J* was initialized with 𝒩 (0, 1*/d*_*N*_). Input and readout are initialized with 𝒩 (0, 1).

To systematically manipulate the underlying selection mechanism, we varied the strength of regularization applied to the recurrent connectivity during training. For each *λ* chosen from the set {0.01, 0.02, …, 0.10}, we trained 10 RNNs. Therefore, in total, 100 RNNs are trained.

#### Details for LINT in the simulated CDM task

For each trained vanilla RNN, we fitted a corresponding low-rank RNN using the LINT framework (Yu et al. 2005). The low-rank model had rank 4 and employed an activation function *ϕ* : *x* → ln(*e*^*x*^ + 1) −1. Each model was trained for 200,000 optimization steps using the Adam optimizer with a learning rate of 0.001 and a batch size of 512. To evaluate model performance and extract the reported results, we generated an independent test set consisting of 5,120 randomly sampled trials.

#### Context-dependent LDS baseline

As a baseline comparison, we fitted a context-dependent linear dynamical system (LDS), following prior deterministic state-space models for neural population activity (Soldado-Magraner et al. 2024). This model extends the standard LDS by allowing the latent dynamics to depend explicitly on task context.

For each context *c*, the latent state evolves according to

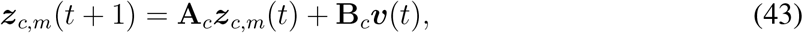

where ***z***_*c,m*_(*t*) ∈ ℝ^*r*^, *r* = 4 denotes the latent state at time *t* for context *c* and stimulus modality *m*, **A**_*c*_ ∈ ℝ^*r*×*r*^ is a context-specific transition matrix, and **B**_*c*_ specifies how external inputs enter the latent dynamics. The input signal ***κ***(*t*) controls the temporal profile of the external drive and is shared across contexts.

The latent state is linearly mapped to the observed neural activity via

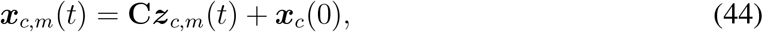

where **C** ∈ ℝ^*N* ×*r*^ is a loading matrix. No explicit observation noise model was assumed; instead, the model was trained deterministically to minimize the reconstruction error of condition-averaged neural activity, consistent with prior deterministic LDS formulations.

Model parameters were optimized by minimizing the mean squared error between model predictions and neural data. After training, the inferred matrices **A**_*c*_ and **B**_*c*_ were used for baseline analyses of context-dependent latent dynamics.

#### Linearized dynamical system analysis for RNNs in the CDM task

To analyze selection mechanisms in RNNs performing context-dependent computation, we followed established linearized dynamical system analyses (Mante et al. 2013). This analysis provides a local linear approximation of the nonlinear RNN dynamics around task-relevant operating points.

The continuous-time dynamics of an RNN in context *c* are given by

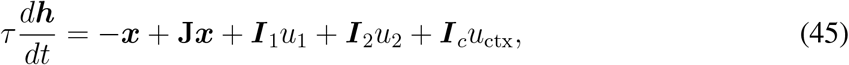

where ***h*** denotes the neural state, ***x*** = *ϕ*(***h***) is the neuronal activity after a nonlinear activation function, **J** is the recurrent connectivity, and ***I***_1_, ***I***_2_, and ***I***_*c*_ denote the input weights for the two sensory modalities and the contextual cue, respectively.

For each network and context, we first identified a slow point ***h***^∗^ using an optimization-based procedure (Mante et al. 2013). At the slow point, the dynamics satisfy

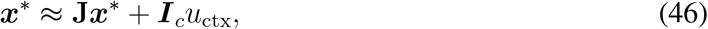

where ***x***^∗^ = *ϕ*(***h***^∗^). We then defined a diagonal gain matrix

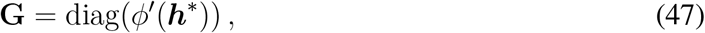

whose diagonal entries capture the local sensitivity of each neuron at the slow point.

Linearizing the dynamics around ***h***^∗^, the evolution of activity perturbations Δ***x*** = ***x*** − ***x***^∗^ can be approximated as

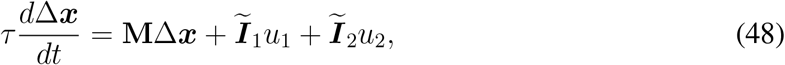

with

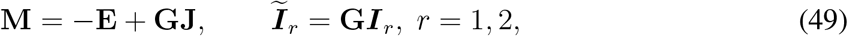

where **E** denotes the identity matrix. Thus, near the slow point, the nonlinear RNN dynamics are approximated by a linear dynamical system.

Consistent with previous work manteContextdependentComputationRecurrent2013, the linearized system typically exhibits a single eigenvalue of **M** close to zero, with all remaining eigenvalues having negative real parts. The right eigenvector associated with the near-zero eigenvalue defines the stable direction of the dynamical systems, called the line-attractor direction ***ρ*** (normalized to unit norm) while the corresponding left eigenvector defines the selection vector ***s***, scaled such that ***s*** · ***ρ*** = 1. Previous studies (Mante et al. 2013) have proved that any perturbation Δ*x*_0_ away from the line attractor will eventually decay back to the line attractor with the distance from the starting point given by Δ*x*_0_ · ***s***.

Following paganIndividualVariabilityNeural2024, we quantified context-dependent computational strategies by decomposing modulation effects into input modulation and selection vector modulation. For a given stimulus input, these quantities are defined as

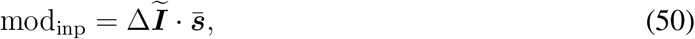

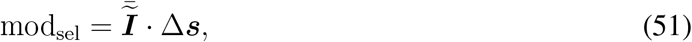

where Δ denotes the difference across contexts and the overbar denotes the average across contexts. The proportion of selection vector modulation is defined as

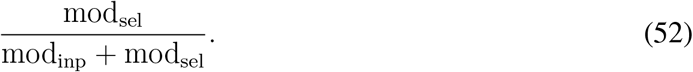

This linearized dynamical system analysis was applied identically to both trained RNNs and LINT-inferred low-rank RNNs.

#### Calculating selection vector for cdLDS

The selection vector in the latent space ***s***_latent_ for cdLDS in context-*c* was defined as the left eigen-vector associated with the near-zero eigenvalue of the latent transition matrix *A*_*c*_. Then the selection vector in the activity space was then obtained by projecting ***s***_latent_ back to the activity space via ***s*** = *C****s***_latent_. The input representation direction was defined as *CB*_*c*_. The input modulation proportion was then defined by Eq. 16.

### Details for fitting PFC recordings in CDM task

#### Experimental setting and neural recording

We analyzed electrophysiological recordings from macaque monkeys performing a context-dependent decision-making (CDM) task (Mante et al. 2013). Each trial began with a 350-ms contextual cue, followed by a 750-ms stimulus period in which a dynamic random-dot display conveyed sensory evidence along two dimensions: motion direction and dot color. Across alternating blocks, the contextual cue instructed the animal to report either the dominant motion direction or the dominant color via a saccadic eye movement.

Neural activity was recorded from the frontal eye field (FEF), a region within the macaque pre-frontal cortex, yielding recordings from 727 neurons in one animal. To obtain smooth population trajectories suitable for dynamical analysis, spike trains during the stimulus period were discretized into 10-ms bins and temporally smoothed using a Gaussian kernel with a standard deviation of 50 ms. For each neuron, trial-averaged firing rates were computed separately for each task condition, resulting in a *T* × 72 data matrix, where *T* = 75 denotes the number of time steps and 72 denotes the number of experimental conditions. Only trials with correct behavioral responses were included in this analysis. To further reduce noise and focus on population-level dynamics, the resulting neural activity was projected onto the low-dimensional subspace spanned by the leading principal components of the population response and then reconstructed in the original neural space, retaining only the components associated with the top 12 principal axes. Finally, the neural responses were demeaned for each neuron at each time step by subtracting the condition-averaged activity, yielding the processed neural activity denoted by *x*_*i,c,t*_, where *i* indexes neurons, *c* indexes task conditions, and *t* = 1, …, 75 indexes time steps within the stimulus period.

#### gmLDS model input setting

The gmLDS model was supplied with four externally specified input channels: two noisy sensory input channels and two contextual cue channels. The sensory inputs corresponded to motion and color evidence, respectively, whereas the contextual cue inputs encoded the task context using binary values (0 or 1). Each trial was modeled as a sequence of 110 discrete time steps. Contextual cue inputs were present throughout the entire sequence, reflecting the persistent contextual instruction, while sensory stimulus inputs were non-zero only during the final 75 time steps, with amplitudes proportional to the coherence of the task-relevant sensory modality.

During training, only neural activity reconstructed during the stimulus period (model time steps *t* = 36, …, 110) was compared against ground-truth neural activity to compute the observation loss term in the evidence lower bound (ELBO). For the initial 35 model time steps (*t* = 1, …, 35), during which only contextual inputs were present and neural recordings were unavailable, neural activity was approximated by a linear ramp,

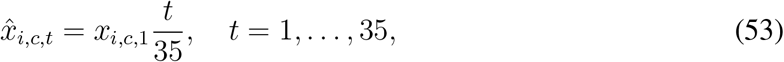

where *x*_*i,c*,1_ denotes the neural activity at stimulus onset. This pre-stimulus padding was introduced to provide a consistent initialization for latent dynamics, being used exclusively by the encoding module and excluded from the reconstruction loss.

#### Targeted dimensionality reduction (TDR)

To visualize trial-averaged population dynamics, we applied targeted dimensionality reduction (TDR) (Mante et al. 2013). At each time point *t*, and for each neuron *i*, we fit a linear regression model to trial-averaged activity:

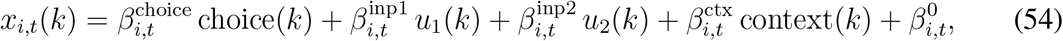

where *x*_*i,t*_(*k*) denotes the activity of neuron *i* at time *t* on trial *k*, and the regressors correspond to choice, the two sensory inputs, and task context.

For each regressor *j*, regression coefficients were pooled across neurons to form a time-varying population regression vector ***β***_*j,t*_. We identified the time point at which each regression vector attained maximal norm,

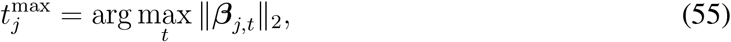

and defined a time-independent regression axis as

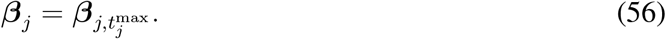

The resulting regression vectors were assembled into a matrix **B** = [***β***_choice_, ***β***_inp1_, ***β***_inp2_, ***β***_ctx_], which was orthogonalized using QR decomposition to obtain an orthonormal TDR basis.

Finally, neuronal activity was averaged across trials sharing the same condition, and the resulting population trajectories were projected onto selected TDR axes (e.g., choice and sensory input axes) to visualize trial-averaged dynamics.

#### gmLDS fitting details

We fit gmLDS models to the processed trial-averaged neural activity. All models were trained using the Adam optimizer with a learning rate of 5 × 10^−4^ and gradient clipping at 0.01, with mini-batches of size 216. Each model was trained for 6.5 × 10^5^ optimization steps steps to ensure convergence of the KL-divergence (Supp. Fig. 6).

We swept the latent dimensionality *d*_*z*_ ∈ {1, 2, 4, 6, 8, 10, 12}. For each *d*_*z*_, a five-fold cross-smoothing scheme was used, with multiple random initializations per fold, and results were aggregated across folds. *α* is set to 0.2 in this model. All analyses in the main text used model of rank-8, as reconstruction performance saturated beyond this value.

### Supplementary figures

## Notes

### Competing Interest Statement

The authors have declared no competing interest.

